# Transgene-free hematopoietic stem and progenitor cells from human induced pluripotent stem cells

**DOI:** 10.1101/177691

**Authors:** Laurence Guyonneau-Harmand, Bruno L’Homme, Brigitte Birebent, Christophe Desterke, Nathalie Chevallier, Loïc Garçon, Hélène Lapillonne, Marc Benderitter, François Delhommeau, Thierry Jaffredo, Alain Chapel, Luc Douay

## Abstract

The successful production of Hematopoietic Stem and Progenitor Cells (HSPCs) from human pluripotent sources is conditioned by transgene delivery^1-5^. We describe here a dedicated and tractable one step, GMP-grade, transgene-free and stroma-free protocol to produce HSPCs from human induced pluripotent stem cells (hiPSCs). This procedure, applied to several sources of hiPSCs with equal efficiency, is based on a directed differentiation with morphogens and cytokines to generate a cell population close to nascent HSPCs or their immediate forerunners i.e., hemogenic endothelial cells^6-9^. Following engraftment into immunocompromised recipients, this cell population was proved capable of a robust myeloid, lymphoid and definitive red blood cell production in sequential recipients for at least 40 weeks. Further identification of the repopulating cells show that they express the G protein–coupled receptor APELIN (APLNR) and the homing receptor CXCR4. This demonstrates that the generation of *bona fide* HSPCs from hiPSCs without transgenes is possible and passes through an early endo-hematopoietic intermediate. This work opens the way to the generation of clinical grade HSPCs for the treatment of hematological diseases and holds promise for the analysis of HSPC development in the human species.

---

While in standard hiPSC differentiation protocols, CD34^+^CD45^+^ hematopoietic cells appeared from bursting Embryoid Bodies (EBs) at day 10 until day 14^1,5,10,11^, here D17 EBs remained compact and spherical without burst therefore assessing a dramatic delay in differentiation (Fig. 1a). These culture conditions, reproducibly applied to three individual hiPSC lines, derived using different reprogramming protocols demonstrated the robustness of the method.

**Fig. 1:**
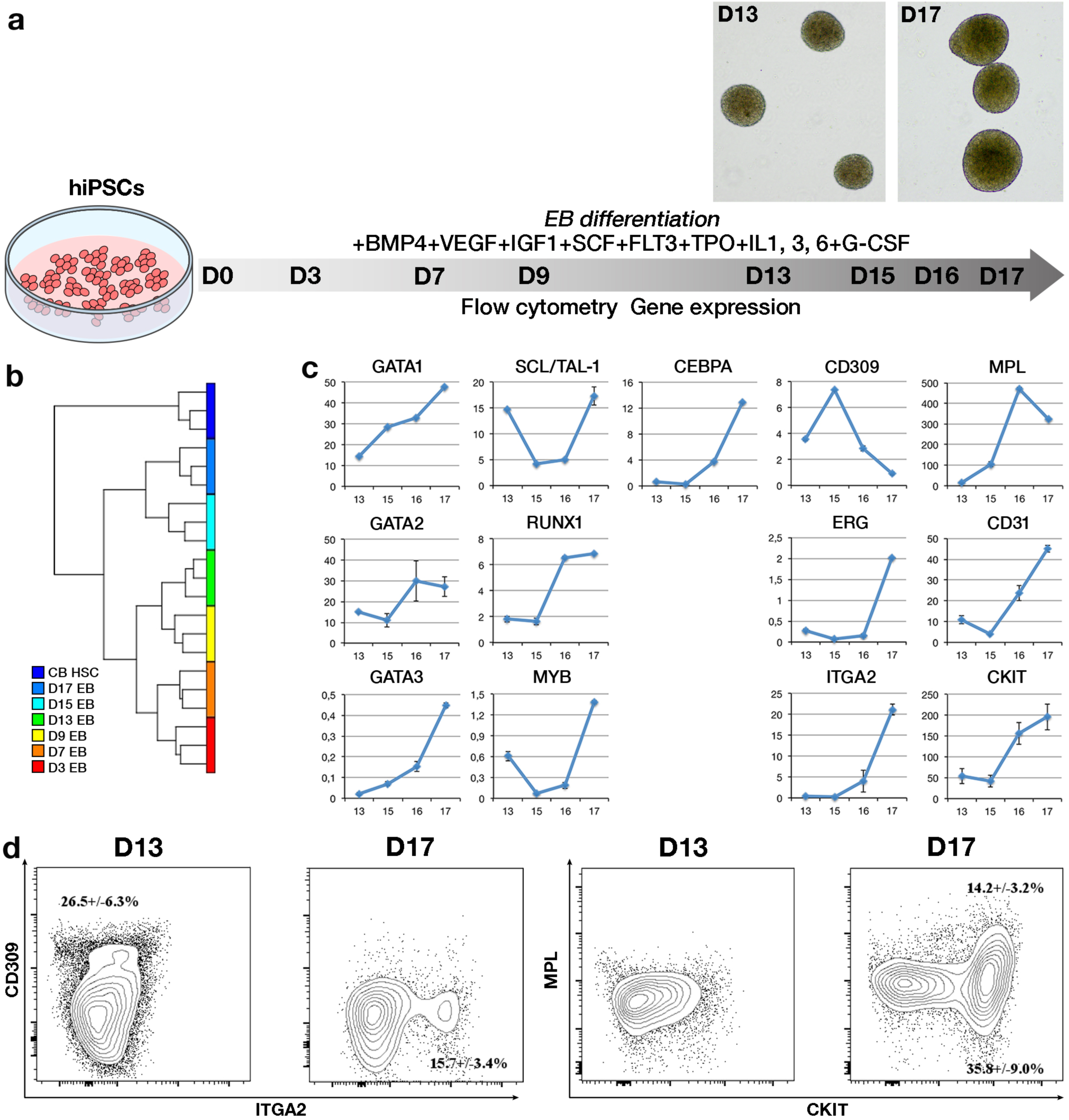
Phenotypic characterization of hiPSC-derived endo-hematopoietic differentiation *ex vivo*. (a) Experimental scheme. HiPSCs were differentiated into EBs over 17 days with the continuous presence of growth factors and cytokines. EBs were characterized at different time points using q-PCR and flow cytometry. Images depicted representative EBs at D13 and 17 respectively. (b) Hierarchical clustering summarizing the expression of the set of 49 genes characteristics of the endothelium, hemogenic endothelium and hematopoietic cells with time in D3, D7, D9, D13, D15 and D17 EBs and in CB CD34^+^ HSPCs. (c) Q-PCR patterns of genes representative of the EHT balance from D13 to D17 EB differentiation. For each gene, the fold change is the mean +/- SEM of 6 experiments (d) Flow cytometry analysis of human CD309, ITGA2, MPL and CKIT expression at D13 and D17 of EB culture.

To determine the point of hemogenic endothelial cell (EC)/early HSPC commitment, we analyzed EBs by qRT-PCR on D3, 7, 9, 13, 15, 16 and 17 for the expression of 49 endothelial- and hematopoietic-specific genes, with CD34^+^ cord blood (CB) HSPCs as a reference (Supplementary Table 1). Hierarchical clustering (Fig. 1b, and Supplementary Fig. 1) revealed two main groups, one associated with CD34^+^ CB HSPCs and another one with D3 to D17 EBs. The latter displayed two distinct clusters: the early EBs (D3 to D13) and the late EBs (D15 to D17). Further analysis of the qRT-PCR identified D13 as the point of EC commitment based on CD309 (VEGFR2/FLK1) mRNA expression and D16 as hemogenic endothelial commitment/EHT based on RUNX1 mRNA expression (Fig. 1c) in keeping with developmental studies^12^. Early HSPC commitment was overt from D17 testified by the expression of ITGA2 (Integrin alpha-2), CEBPA (CCAAT enhancer binding protein alpha), the transcription factors c-MYB and SCL, this latter exhibiting a bimodal curve in keeping with its role on hemangioblast and hematopoietic commitment respectively^13^, the TPO receptor MPL (Fig 1c). Flow cytometry indicated a decrease in CD309 (EC marker) expression from D13 to D17 and an increase of ITGA2, CKIT and MPL (early HSPC markers) expressions (Fig. 1d) in keeping with the qRT-PCR analysis (Fig. 1c).

We next evaluated the endothelial and hematopoietic potential of D15 to D17 EBs using dedicated *ex vivo* functional tests (Fig. 2a and Supplementary Table 2). D15 EBs displayed a strong endothelial-forming potential since they generate endothelial colony-forming cells (CFCs) (Fig. 2a1), pseudo-microtubules (Fig. 2a2) and endothelial-like cells (Fig. 2a3)^14^, but lacked hematopoietic-forming capacity, being unable to generate clonogenic colonies and displaying a low frequency of long term culture-initiating cells (Supplementary Table 2). In contrast D17 EBs lacked endothelial potential but showed an increased hematopoietic capacity (Fig. 2a4, a5), confirming hematopoietic commitment within this period.

**Fig. 2:**
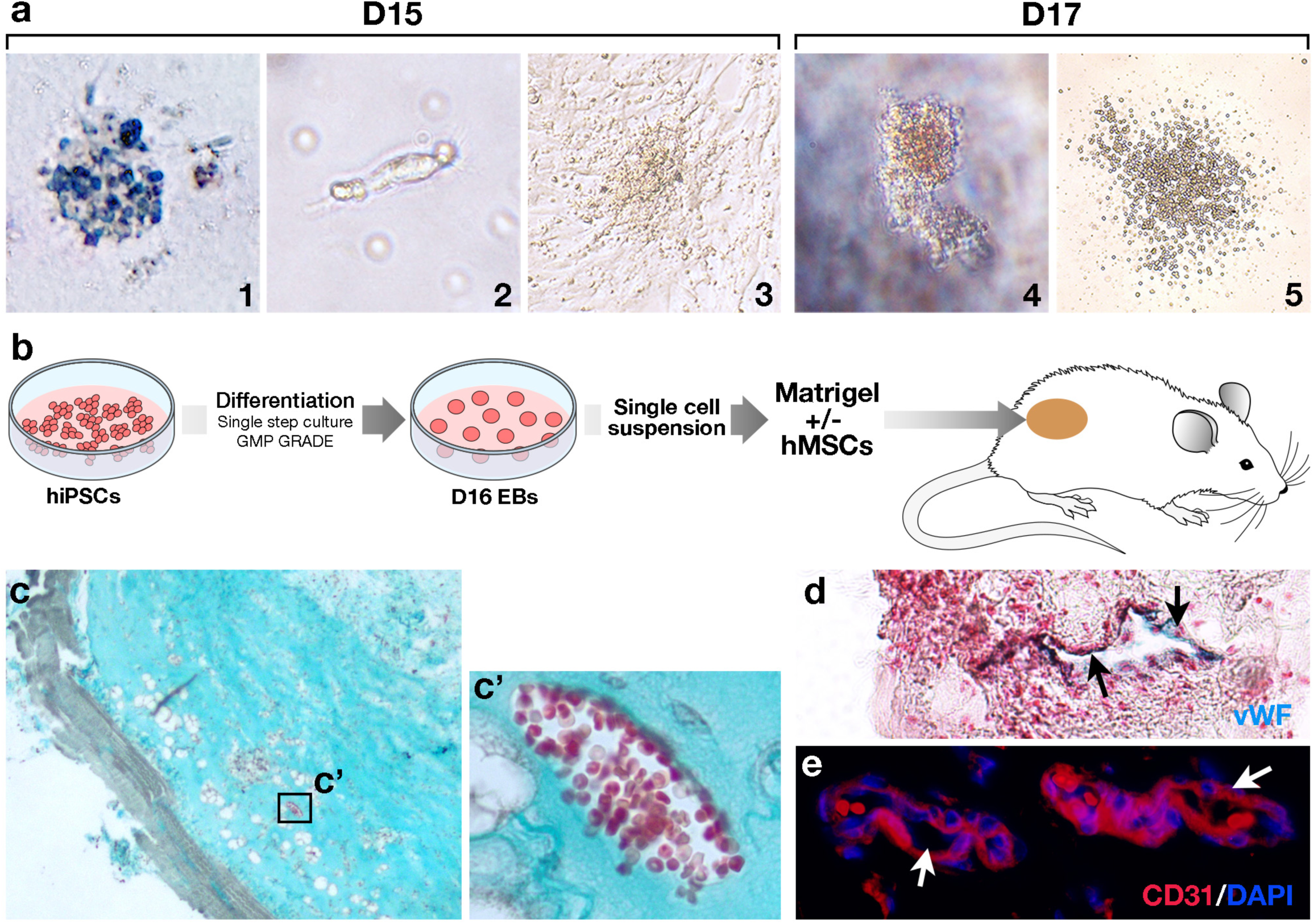
Functional endothelio-hematopoietic profiling between D15 and D17. (a) *Ex vivo* tests probing the presence of endothelial (1-3) and hematopoietic (4-5) progenitors in EBs with time. Dissociated D15-17 EBs generate (1) CFC-ECs, (2) pseudo-microtubules, (3) EC-like cells capable of several passages, (4) CFC, and (5) LTC-ICs. (b) Experimental scheme for *in vivo* tests to probe the endothelial capacity of D16 cells. (c) D16 cells/hMSCs plug section. Masson’s trichrome staining. (d) D16 cells/hMSCs plug section. Human von Willebrand factor^+^ cells (blue) immunostaining. (e) D16 cells/hMSCs plug section. Human CD31^+^ immunostaining (red). Cell nuclei are counterstained by DAPI.

D16 cells were probed *in vivo* by subcutaneous transplantation in Matrigel (growth factor reduced) plug with or without human Mesenchymal Stem Cells (hMSCs) into immunocompromised Foxn1^-/-^ (nude) mice (Fig. 2b)^15^. Two weeks after transplantation, both human vascular and hematopoietic differentiation was found. Human vascular structures (Fig. 2c, d, e), made of human von Willebrand factor^+^ (Fig. 2d) and CD31^+^ (Fig. 2e) cells were detected in the graft seeded with D16 cells associated with (/) hMSCs. qRT-PCR revealed the expression of hVEGFR2, hENG (ENDOGLIN), hPECAM and hVE-CADHERIN in the D16 cells/hMSCs grafts and, as expected, in the endothelial progenitor cells/hMSCs grafts (Supplementary Fig. 2). Moreover, D16 cells/hMSCs grafts also expressed the CD45 antigen and human beta, gamma and epsilon globin transcripts, while D16 cell grafts alone expressed only human epsilon globin transcript disclosing a block of maturation (Supplementary Fig. 2). D16 EBs thus displayed a balanced endothelial-hematopoietic pattern in keeping with our *ex vivo* results.

Based on the newly-formed hematopoietic capability disclosed at D17, we transplanted 4×10^5^ D17 cells intravenously into 30 sublethally irradiated (3.5 gray) 8-week-old immunocompromised mice for 20 weeks followed by a challenging secondary transplantation in similarly-treated immunocompromised recipients for 20 additional weeks and compared it systematically to CD34^+^ CB HSPCs (Fig. 3a, Supplementary Fig. 3a). Hematopoietic reconstitution quantified by the surface expression of hCD45, hCD43 and hCD34 (Fig. 3b and Supplementary Fig. 3b-l)^16^ was evident in 30/30 primary recipients and quantitatively comparable to that of CD34^+^ CB HSPCs (Fig. 3c and Supplementary Fig. 3b, c-e, j) with 20.3+/-2.9 % of hCD45^+^ cells in total mouse BM mononucleated cells, i.e., more than 200 times the threshold of 0.1% usually considered as positive for human hematopoietic engraftment in NSG mice^17^, and 12.2+/-1.5 % of hCD43^+^ and 7.29+/-1.0 % of hCD34^+^. hCD45^+^ BM cells harbor several hematopoietic lineages including B and T lymphoid (hCD19^+^ and hCD3^+^ respectively) and myeloid (hCD14) (Supplementary Fig. 4a-d) whereas hCD45-BM cells harbor hCD235a erythroid progenitors/precursors (Supplementary Fig. 4a-b, e-f). Sorted hCD45^+^ blood cells displayed the same multilineage pattern (Supplementary Fig. 5a) indicating a peripheralization of the grafted cells comparable to results obtained with CD34^+^ CB HSPCs. Their human origin was confirmed (n=30/30) by qRT-PCR using human-specific primers (Supplementary Fig. 4g,). A human-specific clonogenic hematopoietic assay on BM cells from the first recipient revealed an overall frequency of 17.5+/-4.3 clones out of 10^4^ total BM cells (Fig. 3f) distributed into CFU-GEMM, BFU-E and CFU-GM colonies (Fig. 3g1, 2, 3) Cytospin analysis revealed mature macrophages, histiomonocytes, myelocytes and erythroblasts (Fig. 3h1, 2, 3). 7.10^6^ BM cells from the primary recipient were challenged in a secondary (n=30) (Fig 3d, e and Supplementary Fig. 3f-i, k) and eventually a tertiary (n=3) recipient (Supplementary Fig. 3l). Human CD45^+^ cells represented 12.6+/-3.9 % of the mononucleated BM cells at 20 weeks (Fig. 3d), indicating a sustained reconstitution capacity. Multilineage engraftment was found in 30/30 secondary mice (Fig. 3e, Supplementary Fig. 6a-e) and was comparable to the pattern obtained on secondary CD34^+^ CB HSPCs transplants (Supplementary Fig. 3f-i, k and Supplementary Fig. 6). The human CFCs cloning efficiency was 5.5+/-3.1 % in 10^4^ total mouse BM cells, pointing to a robust and prolonged self-renewal potential (Fig. 3f-h). The human origin of the engrafted cells was confirmed as above (Supplementary Fig. 3g).

**Fig. 3:**
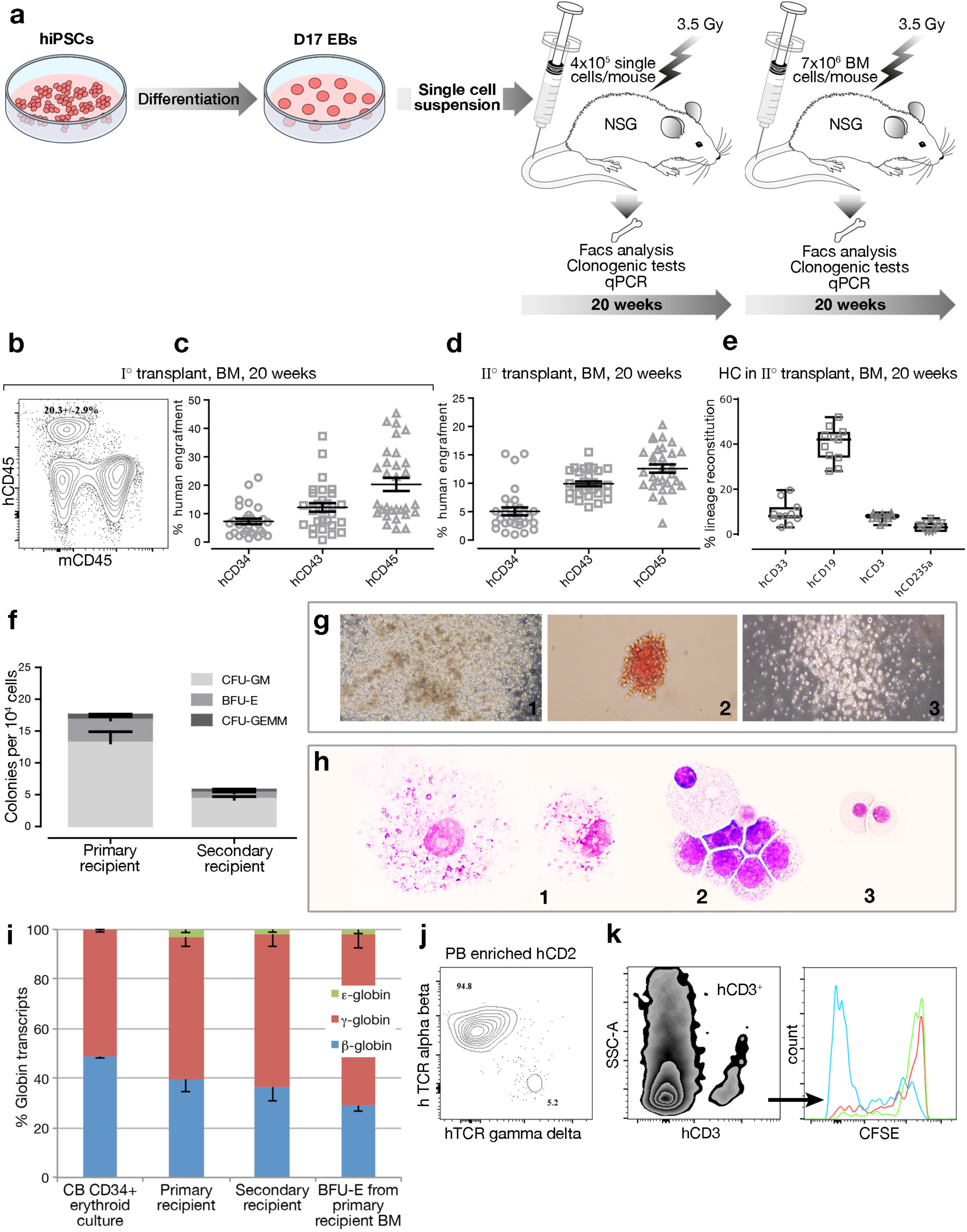
*In vivo* engraftment of D17 EBs in immunocompromised mice. (a) Experimental design. (b) Representative flow cytometry analysis of human vs mouse CD45^+^ cell engraftment in a BM primary recipient. (c-d) Engraftment of hCD34^+^ hCD43^+^ and hCD45^+^ HSPCs in primary (c) or secondary (d) mouse BM 20 weeks post graft. Data are the mean +/- SEM. (e) Human hematopoietic lineage distribution in representative secondary recipients. Cells were gated on hCD45^+^ expression for hCD33, hCD19 and hCD3 whereas hCD235a was analyzed on whole BM cells. Data are the mean +/- SEM. (f) Clonogenic hematopoietic tests on BM cells isolated from primary and secondary recipients. Frequency of CFU-GM, BFU-E and CFU-GEMM colonies. (g) Representative colonies of CFU-GEMM (1), BFU-E (2) and CFU-GM (3) from primary and secondary BM cells. (h) Cytospins. May Grünwald-Giemsa staining of cells isolated from clonogenic tests performed on primary and secondary recipients. Mature macrophages (1), histiomonocytes (2), myelocytes (2) and erythroblasts (3). (i) Human globin expression from CB CD34^+^ HSPCs erythroid culture, from BM primary and secondary recipients and BFU-E from BM primary recipients. Data are mean +/- SEM. (j) Maturation of human T cells. Peripheral blood stained with antibodies against hTCR αβ and hTCR γ δ, the cells are first gated on hCD3 and analyzed for TCR expression. (k) Functionality of human T cells. The whole thymus population is CFSE labeled at D0 and gated on hCD3^+^ expression (green). At D5, the unstimulated population is red while the stimulated parent population is blue.

To ensure the functionality of the grafted cells, we analyzed the ability of the human erythroid precursors from mouse BM to undergo hemoglobin switching *in vivo* and tested the phenotype and the functionality of T cells. Cells from both primary and secondary recipients generated human erythroid progenitors displaying high amounts of β (respectively 39.51+/-4.95% and 36.61+/-5.86%) and γ globin (respectively 57.49+/-3.95% and 61.39+/-4.86%) while ε globin was dramatically reduced to respectively 3.0+/-1.2% and 2.1+/-1.1% of total globin (Fig. 3i), a hallmark of definitive erythrocytes. Peripheral blood hCD3^+^ T cells displayed high amounts of TCRαβ (Fig. 3j) and low amounts of TCRγ δ similar to thymic hCD3^+^ cells (Supplementary Fig. 5b) reminiscent of normal T lymphopoiesis. B and T lymphopoiesis was revealed in spleen CD45^+^ cells by the expression of TCRβ and hCD19 and were found similar to CD34^+^ CB HSPCs grafts (Supplementary Fig 5b). Thymus hCD3^+^ T cells, tested on their ability to expand *ex vivo,* measured by CSFE-labeling, under hCD3 and hCD28 stimulation (Fig. 3k) displayed a high expansion capacity after 5 days, thereby demonstrating T cells functionality.

To further identify the reconstituting population, we focused on APLNR related to ECs^18^ and HCs from human embryonic stem cells^19^ and to the homing receptor CXCR4^20,21^. Differentiating EBs displayed an enhanced APLNR and CXCR4 expression from D7 to D17 with the emergence of a double stained population accounting for 32.1% of the cells at D17 (Fig 4a). Grafting efficiency at 20 weeks was directly proportional to the number of APLNR^+^ cells (Fig 4b). Moreover, only APLNR^+^ cells reconstituted hematopoiesis after 20 weeks with 6.6+/- % hCD45^+^ mononucleated cells, 3.4+/-2.5 % hCD43^+^ and 1.1+/-0.4 % hCD34^+^ in 6/6 grafted mice (Fig. 4c, Supplementary Fig. 7a) and displayed multilineage reconstitution (Supplementary Fig. 7b-c), Importantly, D17 APLNR^+^ cells did not harbor any CD45^+^ cell indicating that reconstitution was not harbored by hCD45^+^ progenitors (Supplementary Fig. 7d). In contrast, APLNR^-^ cells failed to significantly engraft in 4/4 mice with 0.08+/-0.01 % hCD45^+^ cells in mouse BM (Figure 4c, Supplementary Fig 7a). The APLNR^+^ fraction also exhibited a homogeneous population of ENG^+^/TIE^+^/CKIT^+^ (Fig. 4d) described to enhance definitive hematopoiesis in mice^22^. We compared the molecular profiles of APLNR^+^ and APLNR^-^ cells to that of D15 and D17 EBs, to hiPSCs and to control CD34^+^ CB HSPCs through the expression of the set of 49 genes (Supplementary Table 3). Principal component analysis (PCA) of the 49 mRNAs as variables and the six cell populations as observations revealed that the first component likely corresponded to the factor “hematopoietic differentiation” (44.9% of the variance) (Fig. 4e). Aiming to reveal the traits involved in the grafting potential, we compared by PCA the APLNR^+^, D17 and CB HSPC populations to the APLNR^-^ and hiPSC populations. The third component that accounted for 19.23% of the variance segregated two groups differing by their grafting potential (Fig. 4f). A statistical SAM test which measures the strength of the relationship between gene expression and a response variable pointed out 8 genes (FDR<10%) significantly up-regulated in the group unable to graft, among them endothelial genes as TEK, PECAM, and KDR (Fig. 4g).

**Figure 4:**
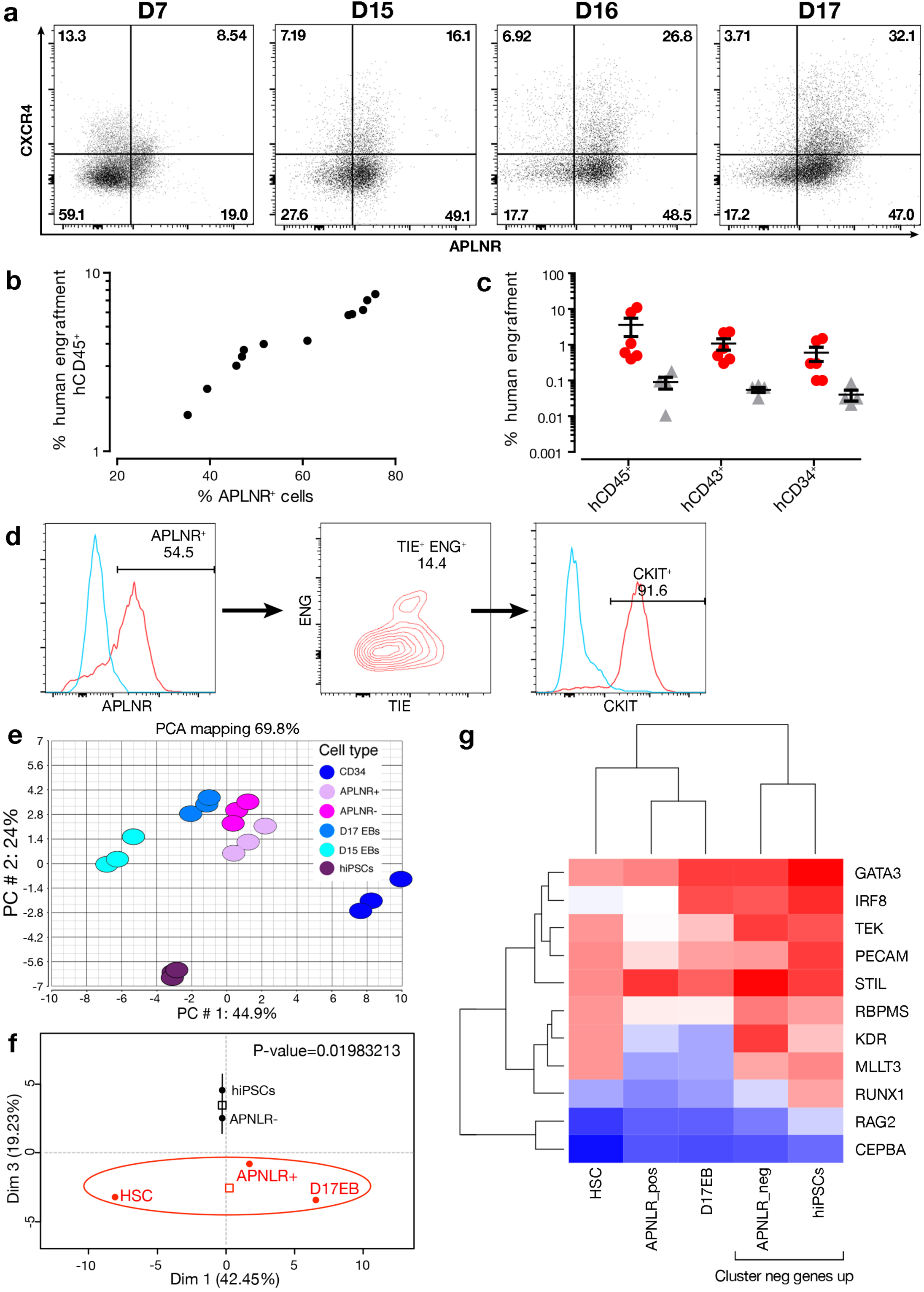
Functional and molecular characterization of the APLNR^+^ population. (a) Flow cytometry analysis of APLNR and CXCR4 expression from D7 to D17 EB culture. (b) Correlation between the percentage of APLNR^+^ cells in the graft to those of hCD45^+^ cells in the NOD-SCID BM primary recipients, 18weeks post-graft. (c) Engraftment capacities of the APNLR^+^ (n=6, red dots) and APNLR^-^ (n=4, grey triangles) populations. Cells containing the reconstitution potential are within the APLNR^+^ population. Data are expressed as the mean +/- SEM percentages of human engraftment, 18 weeks after transplant. (d) Combinatorial flow cytometry analysis of the APLNR^+^ population using CD45, TIE, ENG and CKIT anti-human antibodies. (e) PCA with the set of 49 mRNAs as variables and the six cell populations as observations. PC1 versus PC2 score plot. The PC1 dimension likely corresponds to the trait « hematopoietic differentiation » which accounts for 44.9% of the variance. HiPSCs are segregated from the other populations on the second component which accounts for 24% of the variance. (f) PCA with the set of 49 mRNAs as variables and the populations endowed or not with grafting potential. The PC3 dimension which accounts for 19.23% of the variance segregates the two groups. (g) Heat map of the genes permitting the segregation of the two groups in (f).

Collectively, we show that the generation of long-term multipotent HSPCs supporting hematopoietic reconstitution and self-renewal *in vivo,* passes through an early differentiated cell expressing APLNR more likely an endothelial cell undergoing EHT or a newly formed HSPC. Since this study has been performed under GMP-grade conditions, it may be envisioned that pluripotent stem cells may become a prioritized source of cells for HSPC transplantation.

## Materials and Methods

### hiPSC amplification

The study was conducted using three different hiPSC lines: the FD136-25 (skin primary fibroblasts), reprogrammed with retroviral vectors and Thomson’s combination (endogenous expression of Oct4, Sox2, Nanog and Lin28); the Pci-1426 and Pci-1432 lines (peripheral blood mononuclear cells-Phenocell) reprogrammed with episomes (Sox2, Oct4, KLF, cMyc). hiPSCs were maintained on CellStart (Invitrogen, Carlsbad, USA) in TESR2 medium (Stem Cell Technologies, Bergisch Gladbach, Germany) and the cells were passaged 1:6 onto freshly coated plates every 5 days using standard clump passaging with TRYple select (Invitrogen).

### EB differentiation

EB differentiation was induced as previously described. After 24h, cells were transferred into differentiation medium (Lapillonne et al., 2010) containing 22 ng/mL of SCF, 20 ng/mL of TPO, 300 ng/mL of FLT3, 22 ng/mL of BMP4, 200 ng/mL of VEGF, 50 ng/mL of IL3, 50 ng/mL of IL6, 5 ng/mL of IL1, 100 ng/mL of GCSF, 50 ng/mL of IGF1 (PeproTech, Neuilly-sur-Seine, France). Medium was changed every other day.

### Colony assays

At the indicated times, 1×10^5^ dissociated EBs or 3×10^4^ cells from xenotransplanted recipient BM were plated into 3 mL of complete methylcellulose medium in the presence of SCF, IL-3, EPO and GM-CSF (PeproTech, Neuilly-sur-Seine, France). As G-CSF also stimulates mouse progenitors, it was replaced by granulocyte-macrophage colony-stimulating factor (GM-CSF). Aliquots (1 mL) of the mix were distributed into one 30 mm dish twice and maintained in a humidified chamber for 14 days. Colony-forming Cells (CFC) were scored on day 14.

### Long-term culture-initiating cell assays

Long-term culture-initiating cell (LTC-IC) assays were performed as described previously (*35*), 15–100,000 cells/well on day 17 for the EBs and on day 0 for the control CD34+. Absolute LTC-IC counts corresponded to the cell concentrations, yielding 37% negative wells using Poisson statistics.

### Pseudo-microtubules and EPC-like cells

For Pseudo-microtubules formation, cells were transferred onto growth factor reduced Matrigel (Corning) and culture in EGM2 medium (Lonza).

For EPC-like cells generation, cells were first plated on gelatin and cultured in EBM2 (Lonza) and split several times, after the first passaged the gelatin was no longer mandatory.

### Flow cytometry

Staining of BM cells or dissociated EBs was performed with 2×10^5^ cells in 100 μL staining buffer (PBS containing 2% FBS) with 5:100 dilution of each antibody, for 20 min at room temperature in the dark. Data acquisition was performed on a Becton Dickinson Canto II cytometer.

### In vivo analyses of angiogenesis potential

Foxn1-/-nude mice (Charles River, L’Abresle, France) were housed in the IMRB animal care facility. All experiments and procedures were performed in compliance with the French Ministry of Agriculture regulations for animal experimentation and approved by the local ethics committee.

In vivo assessment of the endothelial and hematopoietic potentials were probed on 9 nude mice. 1.750 10^6^ D16 single cells or hEPCs and 1.750 10^6^ hMSCs were mixed with 100μl of Matrigel phenol red free and growth factor reduced (Corning) and subcutaneously injected into the back of nude mice (two different plugs/ mouse). The controls were performed similarly but with 3.5 10^6^ hMSCs or D16 single cells or hEPCs; for each condition n=3. Two weeks later, mice were sacrificed, the matrigel plug dissected out and cut into two parts. One part was processed for paraffin sectioning. Sections were deparaffinized, hydrated and stained whether with Masson’s trichrome, a three-color protocol comprising nuclear staining with hematoxylin, cytoplasmic staining with acid fuchsin/xykidine ponceau and collagen staining with Light Green SF (all from VWR); or whether with an anti human Von Willebrand factor antibody (Dako), staining was developed with histogreen substrate (Abcys) and counterstained with Fast nuclear red (DakoCytomation), dehydrated and mounted, or wether with hCD31 (R&D system) as primary antibody and donkey anti-rabbit Cy3 (Jackson ImmunoResearch) as secondary antibody and DAPI and mounted with fluoromount G.

### Sorting of APLNR positive cells

Cells were stained with the antibody hAPJ-APC clone 72133 (R&D systems) as described above. Sorting was carried out on a Moflo ASTRIOS Beckman Coulter apparatus and the purity was 98.1% APLNR^+^ cells.

### Mouse transplantation

NOD/SCID-LtSz-scid/scid (NOD/SCID) and NOD.Cg-Prkdc^scid^Il2rg^tm^1^Wjl^/SzJ (NSG) (Charles River, L’Abresle, France) were housed in the IRSN animal care facility. All experiments and procedures were performed in compliance with the French Ministry of Agriculture regulations for animal experimentation and approved by the local ethics committee.

Mice, 6-8 weeks old and raised under sterile conditions, were sublethally irradiated with 3.5 Grays from a 137Cs source (2.115 Gy/min) 24 h before cell injection. To ensure consistency between experiments, only male mice were used. Prior to transplantation, the mice were temporarily sedated with an intraperitoneal injection of ketamine and xylazine. Cells (4 × 10^5^ per mouse) were transplanted by retro-orbital injection in a volume of 100 μL using a 28.5 gauge insulin needle. A total of 147 mice were used in this study.

For the engraftment potential of the D17 cells on the three different hiPSC lines:

86 NSG mice were used as followed: 30 primary recipients, 30 secondary recipients and 26 as control.

48 NOD-SCID mice were used as followed: 20 primary recipients and 16 secondary recipients, 3 tertiary recipients and 9 as control.

For the engraftment potential of APLNR^+^ and APLNR^-^ population: 10 NOD-SCID mice were used and 3 NOD-SCID as control.

### Assessment of human cell engraftment

Mice were sacrificed at week 12, 18 or 20. Femurs, tibias, liver, spleen and thymus were removed. Single cell suspensions were prepared by standard flushing and aliquots containing 1×10^6^ cells were stained in a total volume of 200μL staining buffer.

Samples were stained for engraftment assessment with the following markers: hCD45 clone J33, hCD43 clone DFT1, hCD34 clone 581 (Beckman Coulter) and hCD45 clone 5B1, mCD45 clone 30F11 (Miltenyi)

The BM of three mice were pooled to allow hCD45 microbead enrichment (Miltenyi), the multilineage was assessed using the following human markers: hCD3 clone UCHT1, hCD4 clone 13B8.2, hCD8 clone B9.11, hCD14 clone RMO52, hCD15 clone 80H5, hCD19 clone J3-119, hCD20 clone B9E9, hCD43 clone DFT1, hCD34-APC, hCD71 clone YDJ1.2.2 (all from Beckman Coulter antibodies, Brea, USA), CD45 clone 5B1 (Miltenyi), CD235a clone GA-R2 (Becton-Dickinson).

The blood samples of three mice were pooled to allow hCD45 microbead sorting (Miltenyi). The multilineage potential was assessed using the following human markers: hCD3 clone UCHT1, hCD4 clone 13B8.2, hCD8 clone B9.11, hCD14 clone RMO52, hCD15 clone 80H5, hCD11B Bear1, hCD19 clone J3-119, hCD20 clone B9E9, hIGM clone SA-DA4 (all from Beckman Coulter antibodies, Brea, USA).

Non-injected mouse BM was used as a control for non-specific staining.

Compensation was performed by the FMO method with anti-mouse Ig and data were acquired on a BD Canto II cytometer.

### T-cell maturity and functionality assay

The presence of TCR αβ and TCR γ δ in peripheral blood was assessed by flow cytometry using the following human markers: hCD3 clone UCHT1 (positive gating), TCR αβ clone IP26A and TCR γ δ clone IMMU510 (all from Beckman Coulter antibodies, Brea, USA).

Thymus and spleen cells were isolated, CFSE labelled and seeded in cell culture media complemented or not with hCD3 and hCD28 (Beckman Coulter both 1μg/ml). After 5 days, cells were harvested and stained with anti-hCD3 clone UCHT1 and analyzed on a BD Canto II cytometer. FlowJo analysis software was used to gate on CD3^+^ T-cells and generate the overlaid histogram plots.

### Assessment of the APLNR cell safety

Three sub-lethally irradiated NOD/SCID mice were subcutaneously injected each with 3 million APLNR positive cells. No teratoma was found after 2 months follow-up according to FDA guidelines.

In addition, no tumor was macroscopically detected in any mouse after analysis of the organs (140/140 mice) or after microscopic analysis of different tissues (brain, lungs, kidneys, BM, liver and gut) (140/140 mice).

### Quantitative PCR

Total mRNA was isolated with the RNA minikit (Qiagen, Courtaboeuf, France). mRNA integrity was checked on a Bioanalyzer 2100 (Agilent Technologies, Massy, France). cDNAs were constructed by reverse transcription with Superscript (Life Technologies, Carlsbad, USA). PCR assays were performed using a TaqMan PCR master mix (Life Technologies) and specific primers (Applied BioSystems, Carlsbad, USA) for selected genes (see table below), together with a sequence detection system (QuantStudio(tm) 12K Flex Real-Time PCR System, Life Technologies). In each sample the fluorescent PCR signal of each target gene was normalized to the fluorescent signal of the housekeeping gene glyceraldehyde 3-phosphate dehydrogenase (GAPDH).

The human origin of the mRNAs from mouse BM was assessed by measuring hCD45, hCD15, hMPO, hITGA2 and hGAPDH. From CFCs post grafting and globin type expression in the mouse BM, we measured beta, gamma and epsilon globins using Taqman probes.

Controls were cultured erythroblasts generated from cord blood CD34^+^.

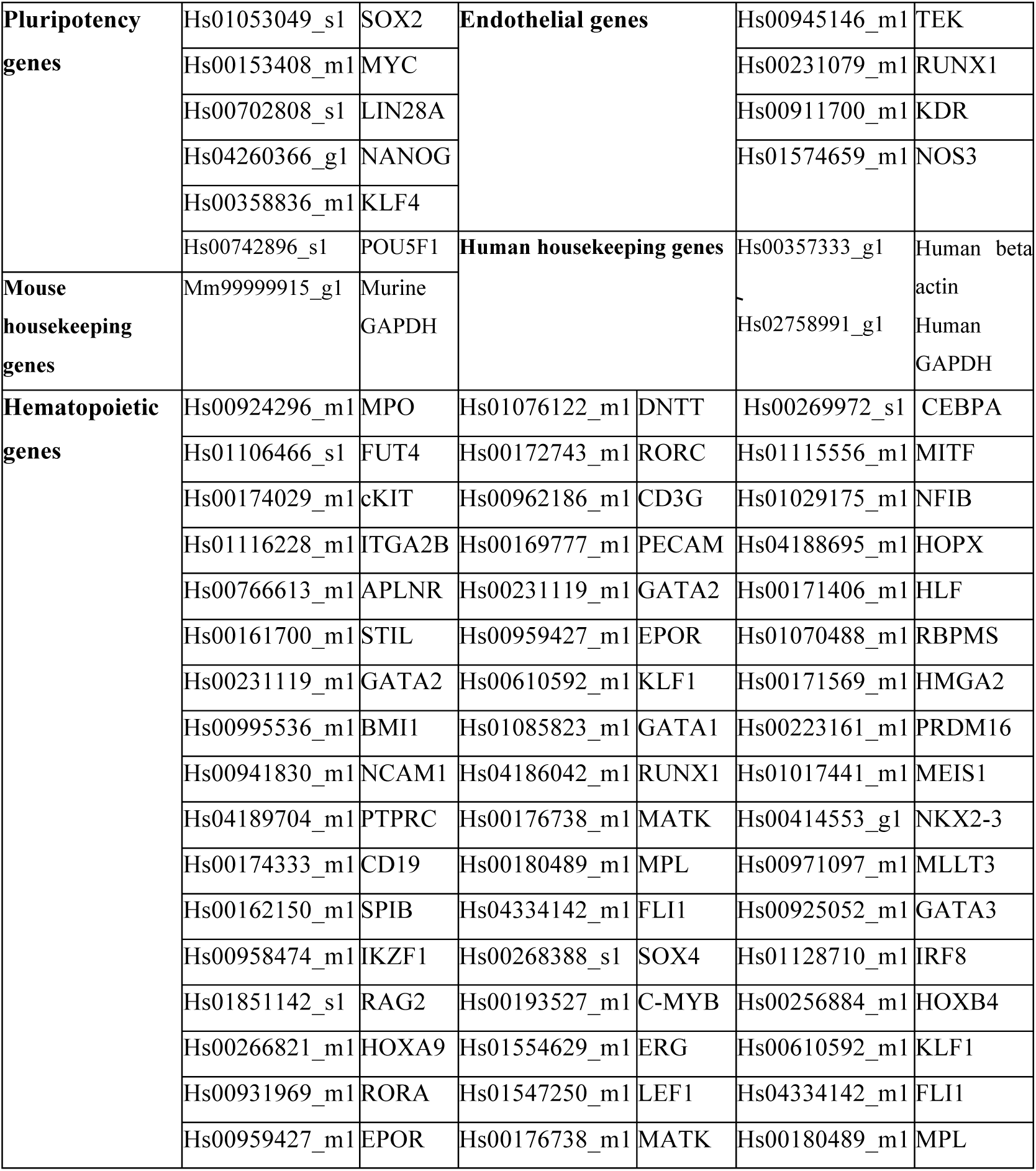

### Statistical analysis

All statistics were determined with R Software 3.1.1 (2014-07-10) (R Core Team, 2013), INGENUITY and SAM Software. Data are represented with hierarchical clustering and PCA.

### References

H. Lapillonne *et al.*, Red blood cell generation from human induced pluripotent stem cells: perspectives for transfusion medicine. Haematologica. 95:1651-1659 (2010).

## Aknowledgments

We would like to thank Franck Chiappini for help in statistical analysis, Charles Durand for expert assistance with heat map and PCA analysis, Stéphane Viville for providing iPS line, Georges Tarlet, Christine Linard, Marianne Gervais-Taurel and Dhouha Darghouth for their technical support, Rima Haddad for her scientific support and Sophie Gournet for excellent drawing assistance. This work was supported by “Direction Générale de l’Armement” via the ASTRID/ANR program, the Etablissement Français du Sang (EFS) via the APR 2013 and the association “Combattre La Leucémie”. These studies were supported by a joint grant from Agence Nationale pour la Recherche/California Institute for Regenerative Medicine (ANR/CIRM 0001-02) for TJ.

## Author contributions

L.G.H. and L.D. designed the study, analyzed the data, and wrote the manuscript. A.C. designed the study. L.G.H. performed experiments with assistance of B.L., B.B. and N.C.; H.L., C.D. and L.G. helped to analyze the results; T.J. analyzed the data and wrote the manuscript. M.B. and F.D. helped design the study; A.C. and L.D. supervised the study.

## Supplementary information

### Inventory of the supplementary figures and tables

#### Supplementary figure 1 is related to Figure 1

Shows the heat map of the gene list and the associated hierarchical clustering. It shows that CD34^+^ cells form a first main branch and that the EBs are segregated in a second main branch. D3 to D13 EBs are separated from 1D5 to D17 EBs.

#### Supplementary figure 2 is related to Figure 2

Endothelial and hematopoietic potential carried by D16 EBs *in vivo*.

#### Supplementary figure 3 is related to Figure 3

It gives a complete overview of the analysis of the hematopoietic engraftment with representative examples of multilineage reconstitution.

#### Supplementary figure 4 is related to Figure 3

Bone marrow analysis of the hematopoietic engraftment, multilineage reconstitution of primary recipients.

#### Supplementary figure 5 is related to Figure 3

A complete overview of the hematopoietic multilineage engraftment of peripheral blood, thymus and spleen from D17 EB cell and CB CD34^+^ HSPCs.

#### Supplementary figure 6 is related to Figure 3

Bone marrow analysis of the hematopoietic engraftment, multilineage reconstitution of secondary recipients.

#### Supplementary figure 7 is related to Figure 4

Characteristics of the APLNR^+^ population in terms of hematopoietic engraftment.

#### Supplementary table 1 is related to Figure 1

*Ex vivo* endothelial and hematopoietic potential carried by hiPSC, HSC and D3 to D17 EBs.

#### Supplementary table 2 is related to Figure 2

*Ex vivo* endothelial and hematopoietic potential of EBs from D15 to D17.

#### Supplementary table 3 is related to Figure 4

*Ex vivo* endothelial and hematopoietic potential carried by hiPSC, CD34+ CB HSPC and APLNR+, APLNR- and D17 EBs.

### Supplementary Legends

**Supplementary Figure. 1.**
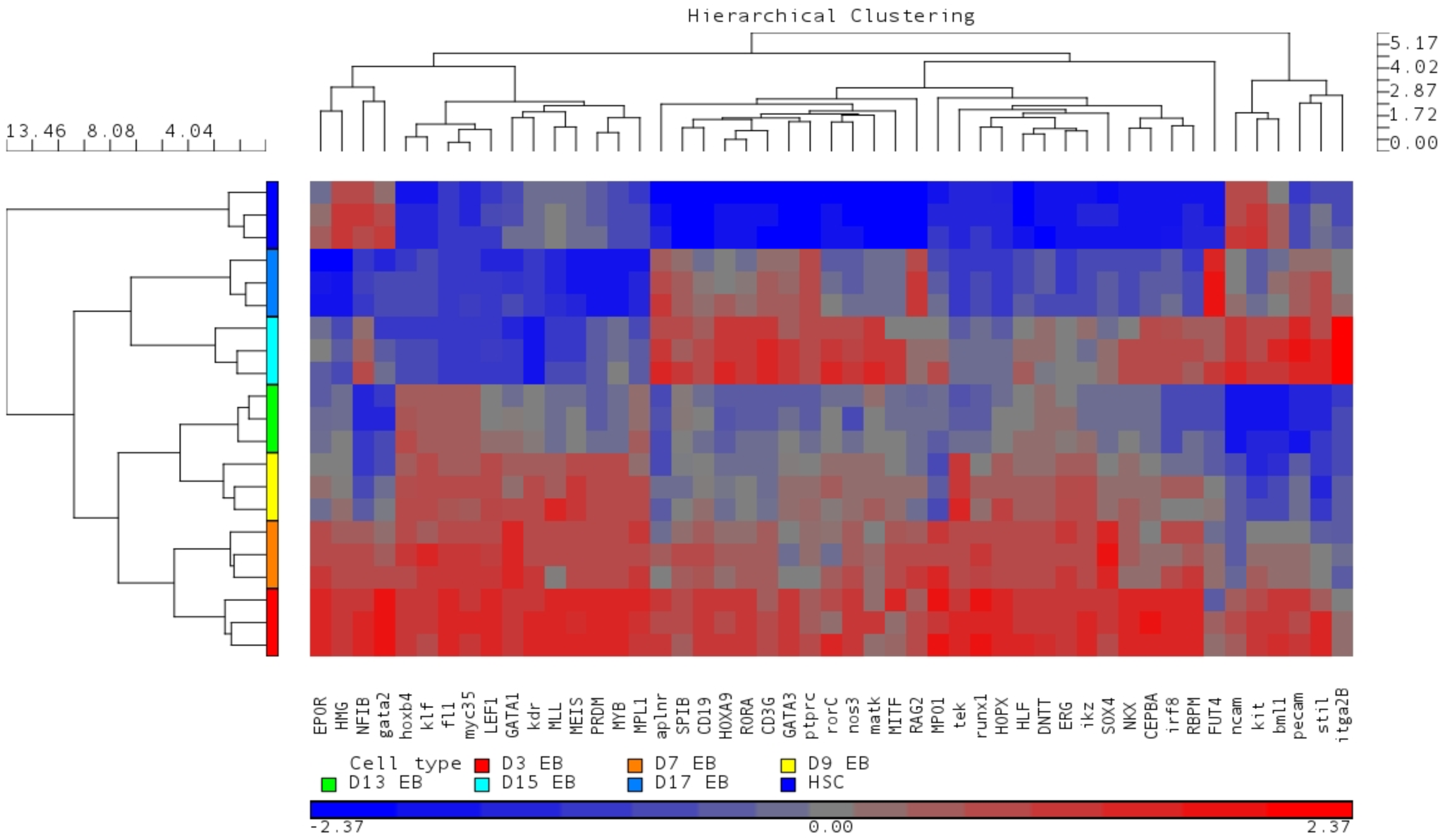
Hierarchical clustering and principal component analysis of the set of 49 genes representative of endothelial and hematopoietic commitment. Heat map of the gene expression and the associated hierarchical clustering. CD34^+^ CB HSPCs are segregated from the EBs. D3 to D13 EBs are separated from D15 to D17 EBs.

**Supplementary Figure. 2.**
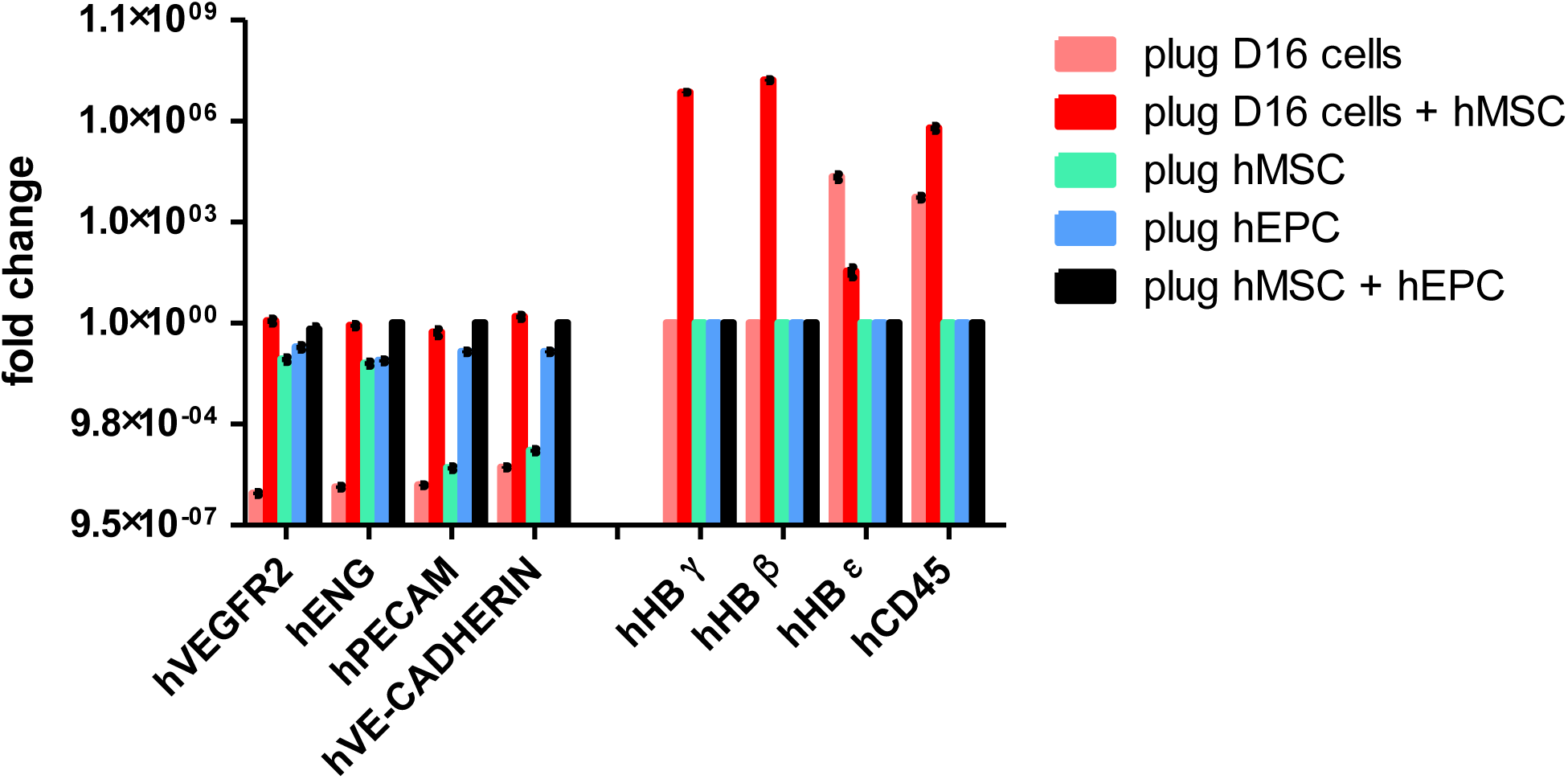
Endothelial and hematopoietic potential carried by D16 EBs *in vivo*. Levels of expression of hVEGFR2, ENDOGLIN, PECAM-1, VE-CADHERIN, γ-globin, β-globin, e-globin and CD45 in the matrigel plugs. For each gene, the fold change is the mean ± SEM of 3 independent experiments.

**Supplementary Figure. 3.**
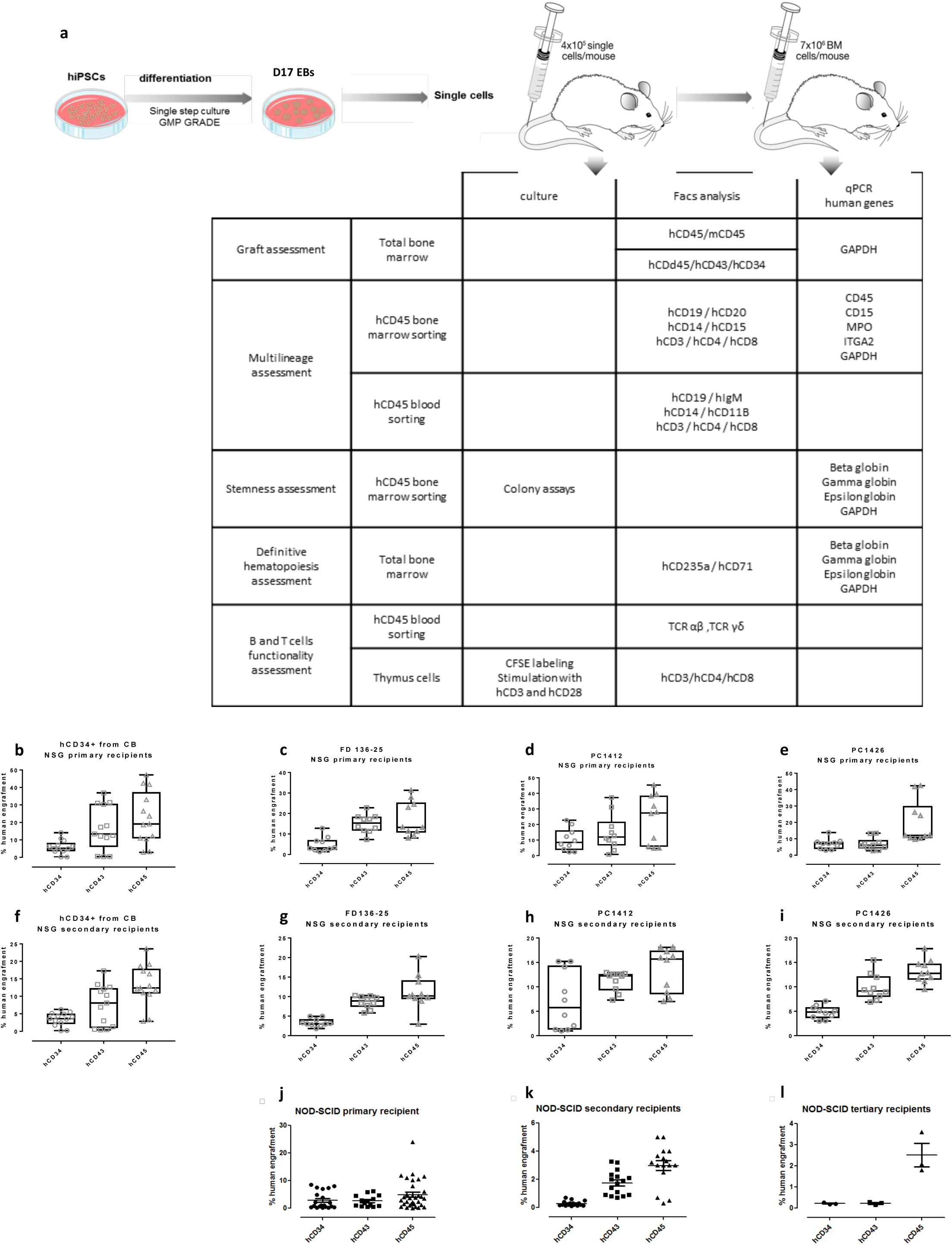
*In vivo* engraftment of D17 human EBs in immunocompromised mice. (a) Experimental design. The table summarizes the different tests and analyses performed on the hematopoietic populations. (b-l) *In vivo* engraftment capacity in immunocompromised mice. (b) CD34^+^ CB HSPCs in NSG primary recipient (n=13) at 20 weeks post-graft after inoculation of 2×10^5^ CD34+ cells. results are in % of CD34^+^, CD43^+^ and CD45^+^ human cells. (c-e) D17 EBs from the three different hiPSCs lines (FD 136-25, PC1412 and PC1425) in NSG primary recipient (n=30) at 20 weeks post-graft after inoculation of 4×10^5^ D17 EBs. Results are in % of CD34^+^, CD43^+^ and CD45^+^ human cells. (f-i) Follow up of the secondary recipients (f) NSG secondary recipient at 20 weeks post-graft after inoculation of 7×10^6^ total BM cells from b. (g-i) NSG secondary recipient at 20 weeks post-graft after inoculation of 7×10^6^ total BM cells from respectively c, d, e. (j-l) Follow up of D17 EBs from the FD136-25 hiPSC line grafted in primary (e, 4×10^5^ cells, n=20), secondary (f, 7×10^6^ total BM cells, n=16) and tertiary (g, 7×10^6^ total BM cells, n=3,) NOD-SCID recipients. Results are in % of human CD34^+^ CD43^+^ CD45^+^ cells in mouse BM at 20 (e), 20 (f) and 12 (g) weeks after transplant. Data are mean +/- SEM.

**Supplementary Figure. 4.**
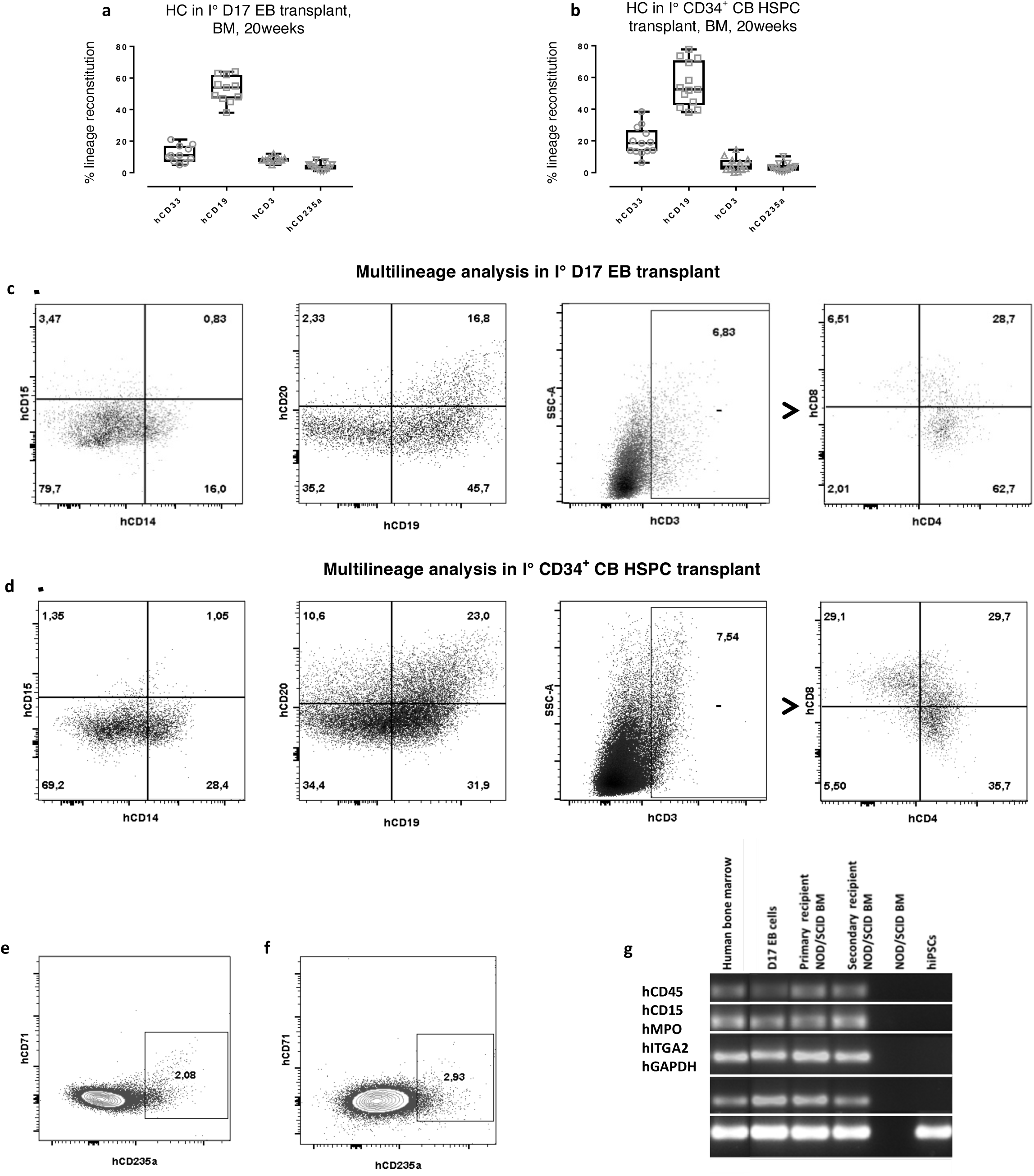
Flow cytometry and qRT-PCR analysis of hematopoietic engraftment in primary BM recipients. (a-b) Human hematopoietic cell lineages in primary recipients: (a) grafted with D17 EBs (n=30) and (b) grafted with CD34^+^ CB HSPCs (n=13). Cells were gated on hCD45+ expression for hCD33, hCD19 and hCD3 whereas hCD235a was analysed on whole BM cells. Data are mean +/- SEM. (c-d) Flow cytometry analysis for myeloid (hCD14/hCD15) and lymphoid (hCD19/hCD20, SSC-A/hCD3 and hCD4/hCD8) lineages from a representative NSG primary recipient grafted with D17 EB cells (c) or with CD34^+^ CB HSPCs (d). Analyses were performed on hCD45^+^ gated BM cells. (e-f) Flow cytometry analysis of the erythroid lineage from a representative NSG primary recipient grafted with D17 EBs (e) or with CD34^+^ CB HSPCs (f). Analyses were performed on whole BM cells. (g) qRT-PCR analysis for hCD45, hCD15, hMPO, hITGA2 and hGAPDH expression in human BM cells (positive control), D17 EBs, NOD/SCID BM primary recipient grafted with D17 EBs, NOD/SCID BM secondary recipient, NOD/SCID BM ungrafted control, FD136-25 hiPSCs. PCR samples were run on several agarose gels. For presentation purpose, bands were assembled one above the other (white lines).

**Supplementary Figure. 5.**
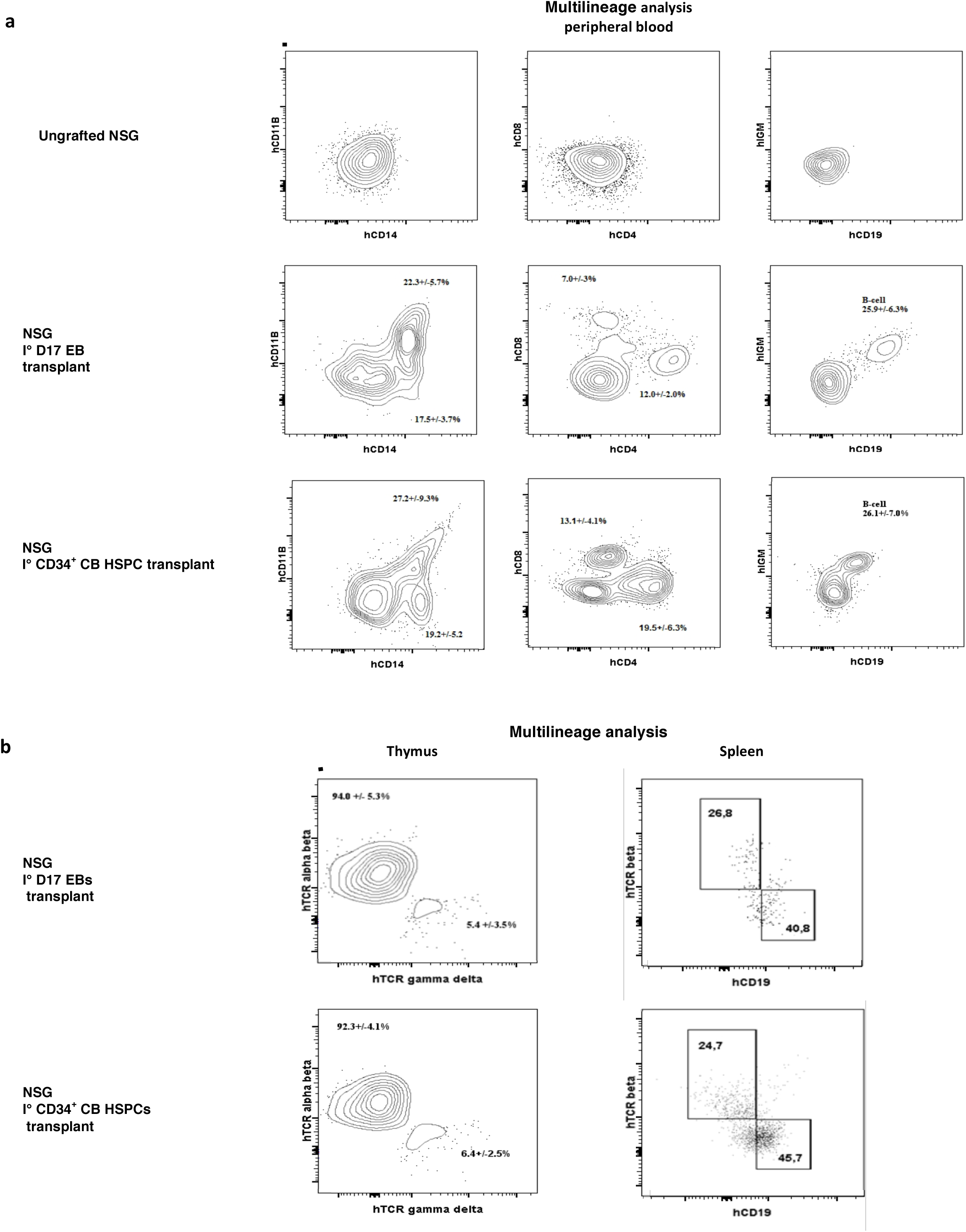
Flow cytometry analysis of human hematopoietic lineages from representative primary recipients peripheral blood, thymus and spleen. (a) Myeloid (hCD11B/hCD14) and lymphoid (hCD19/hIGM and hCD4/hCD8) lineages on peripheral blood of ungrafted recipient (1^st^ line), of D17 EBs grafted recipient (2^nd^ line) and of CD34^+^ CB HSPCs grafted recipient (3^rd^ line). The analyses were performed on hCD45^+^ gated cells, aside the ungrafted recipient. (b) Representative flow cytometry analysis of thymus and spleen cells. Thymus hCD3 gated cells exhibited a large amount of TCR alpha beta cells very close to that obtained using CD34^+^ CB HSPCs. hCD45^+^ gated spleen cells exhibited B or T lymphoid phenotype with ratios similar to those obtained with CD34^+^ CB HSPCs.

**Supplementary Figure. 6.**
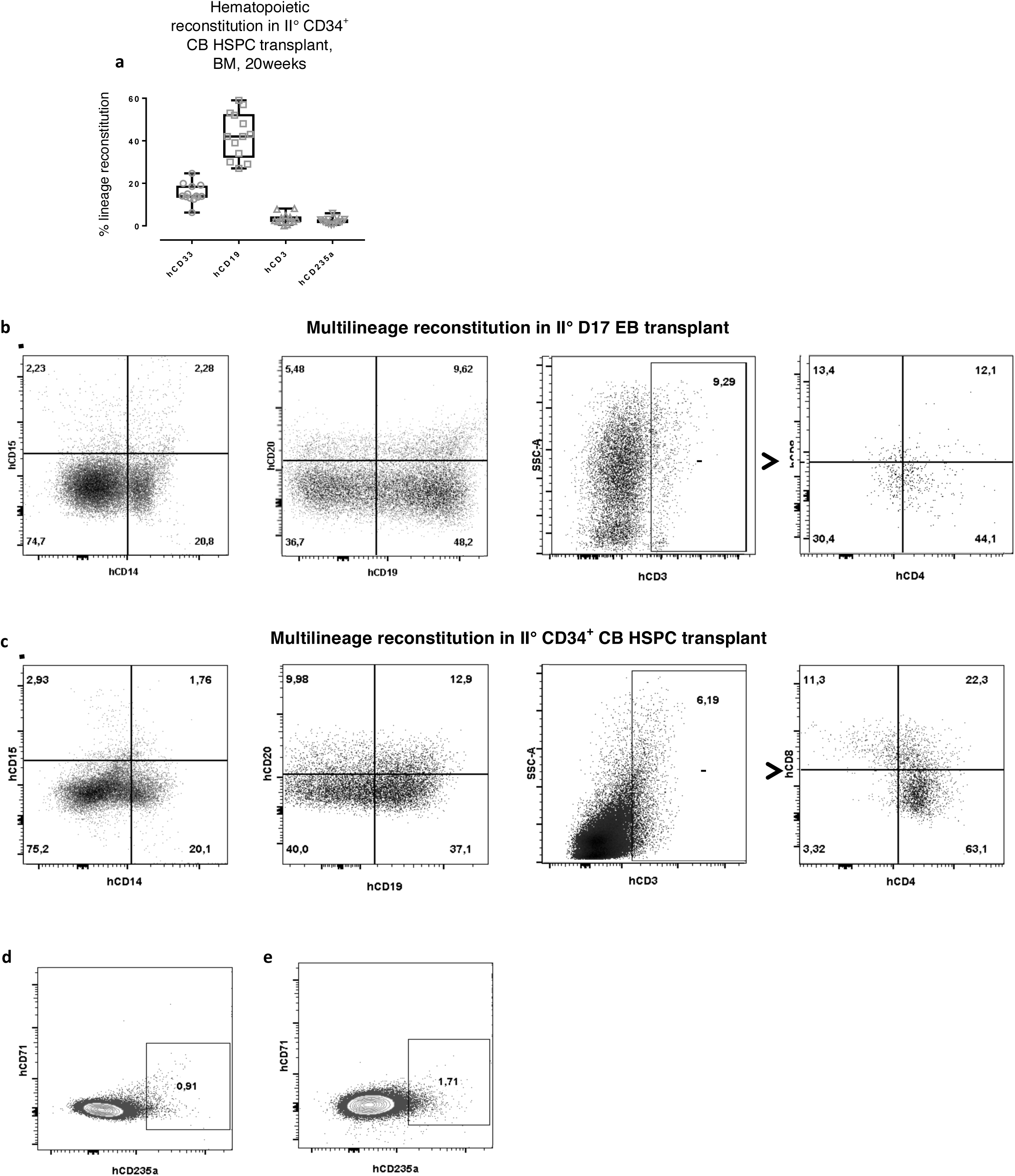
Flow cytometry analysis of hematopoietic engraftment in secondary BM recipients. (a) Human hematopoietic lineage distribution in secondary recipients grafted with 7.10^6^ whole bone marrow from CD34^+^ CB HSPCs-grafted primary recipient (n=13). Cells were gated on hCD45+ expression for hCD33, hCD19 and hCD3 whereas hCD235a was analysed on whole BM cells. Data are mean +/- SEM. (b-c) Flow cytometry analysis of the myeloid (hCD14/hCD15) and lymphoid (hCD19/hCD20, SSC-A/hCD3 and hCD4/hCD8) lineages. (b) From a representative NSG secondary recipient grafted with 7.10^6^ whole BM from D17 EB primary recipient. Analyses were performed on hCD45^+^ gated BM cells. (c) From a representative NSG secondary recipient grafted with 7.10^6^ whole BM from CD34^+^ CB HSPCs-grafted primary recipient. (d-e) Representative flow cytometry analysis of the erythroid lineage. Analyses performed on whole BM cells.: (d) Representative NSG secondary recipient grafted with 7.10^6^ whole BM of D17 EBs primary recipient and (e) Representative NSG secondary recipient grafted with 7.10^6^ whole BM CD34^+^ CB HSPCs-grafted primary recipient.

**Supplementary Figure. 7.**
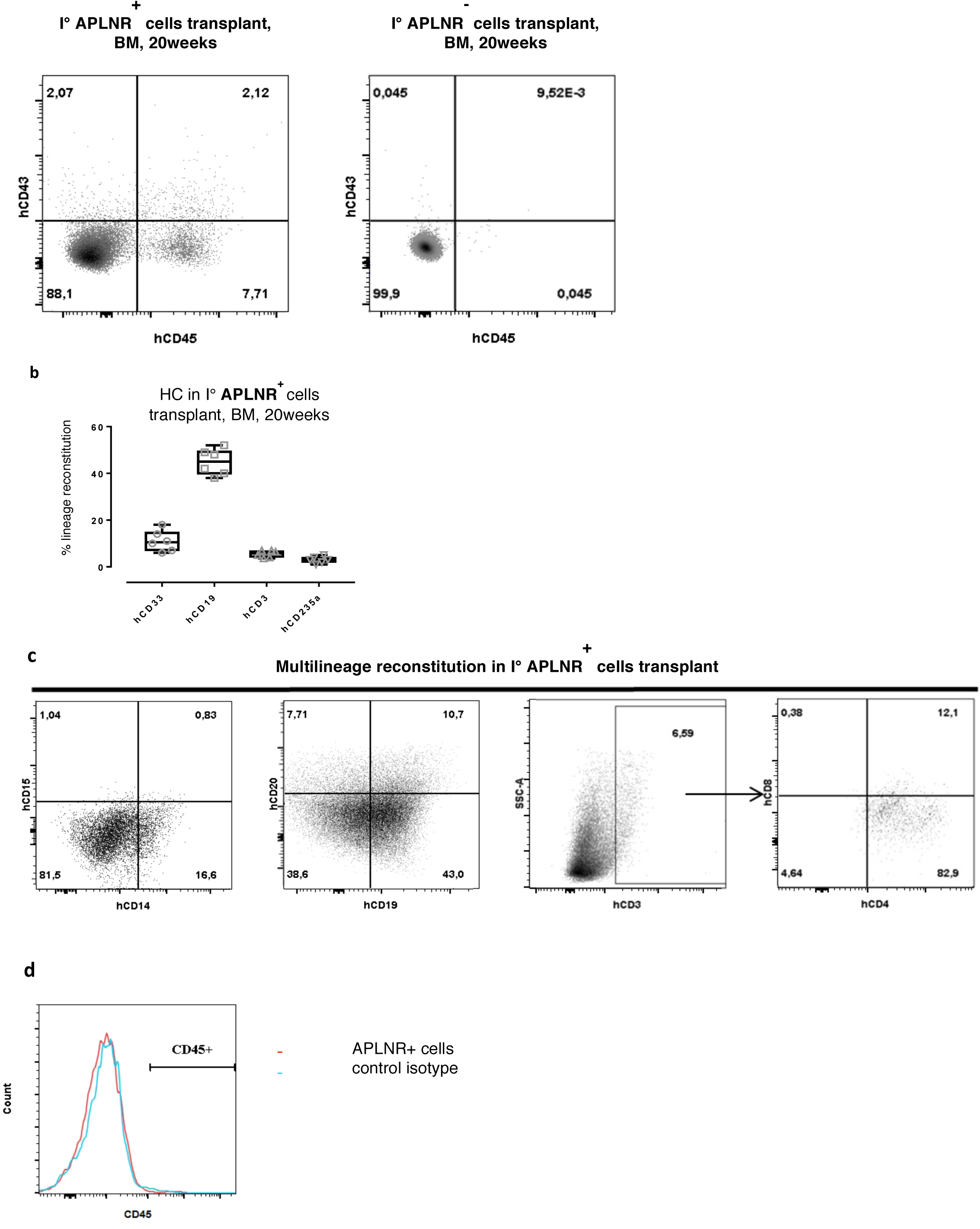
Characterization of the APLNR^+^ and APLNR ^-^ populations. (a) Representative profile of hCD45 and hCD43 expression in mice BM grafted with either the D17 APLNR^+^ (left) or ^−^ (right) cell fractions. (b) Human hematopoietic lineage distribution in primary BM recipient grafted with APLNR^+^ cells (n=6). Data are mean +/- SEM. (c) Representative flow cytometry analysis of the myeloid (hCD14/hCD15) and lymphoid (hCD19/hCD20, SSC-A/hCD3 and hCD4/hCD8) lineages present in a primary BM recipient grafted with APLNR^+^ cells. (d) Representative flow cytometry analysis showing the absence of CD45 expression on APLNR^+^ cells

### Supplementary Tables

**Supplementary Table 1. Ex viv*o* endothelial and hematopoietic potential carried by hiPSC, CD34^+^ CB HSPC and D3 to D17 EBs.**

Refer to the attached supptable1.xlsx file.

**Supplementary Table 2.**
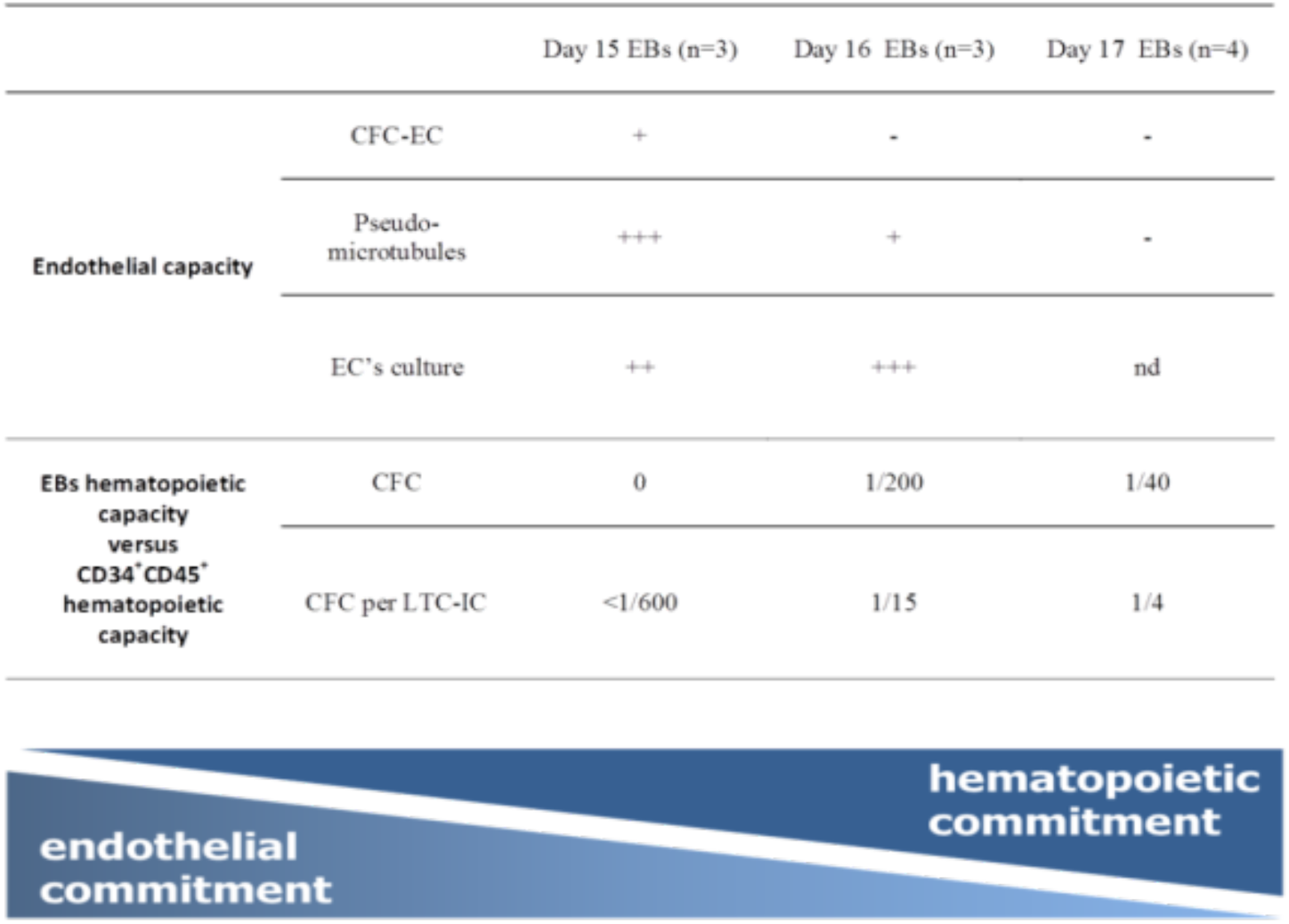
Endothelial and hematopoietic potential carried by D15 to D17 EBs *ex vivo.* EBs were probed for their capacities to give rise to either EC or HC colonies based on recognized functional tests. The highest endothelial potential was found at D15, markedly decreased at D16 and was absent at D17. In contrast, the hematopoietic potential was first detected at D16 and was high at D17. Legend: EB: embryonic bodies; LTC-IC: Long Term Culture-Initiating Cells; CFC: Colony-Forming Cell.

**Supplementary Table 3.**
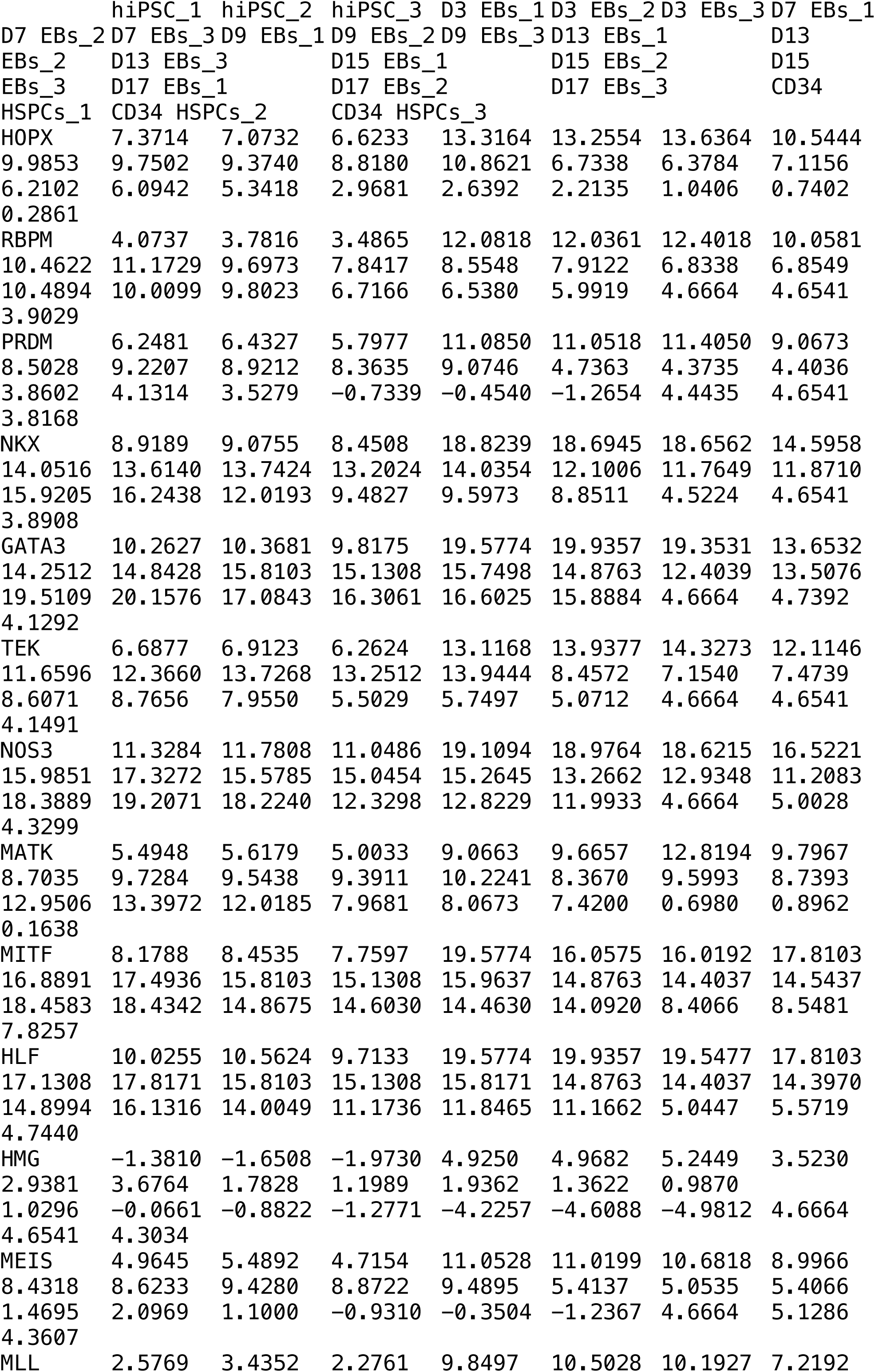
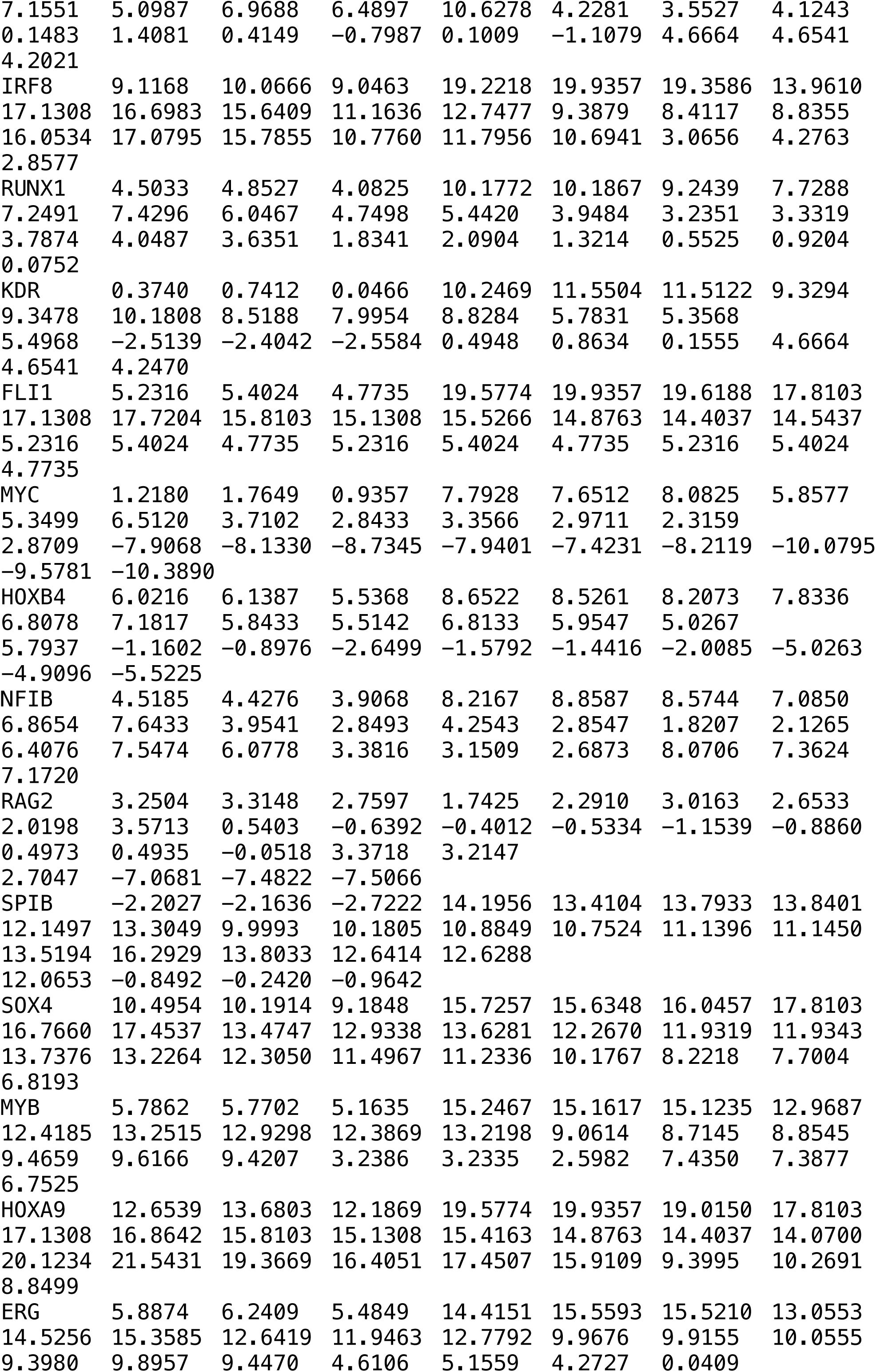
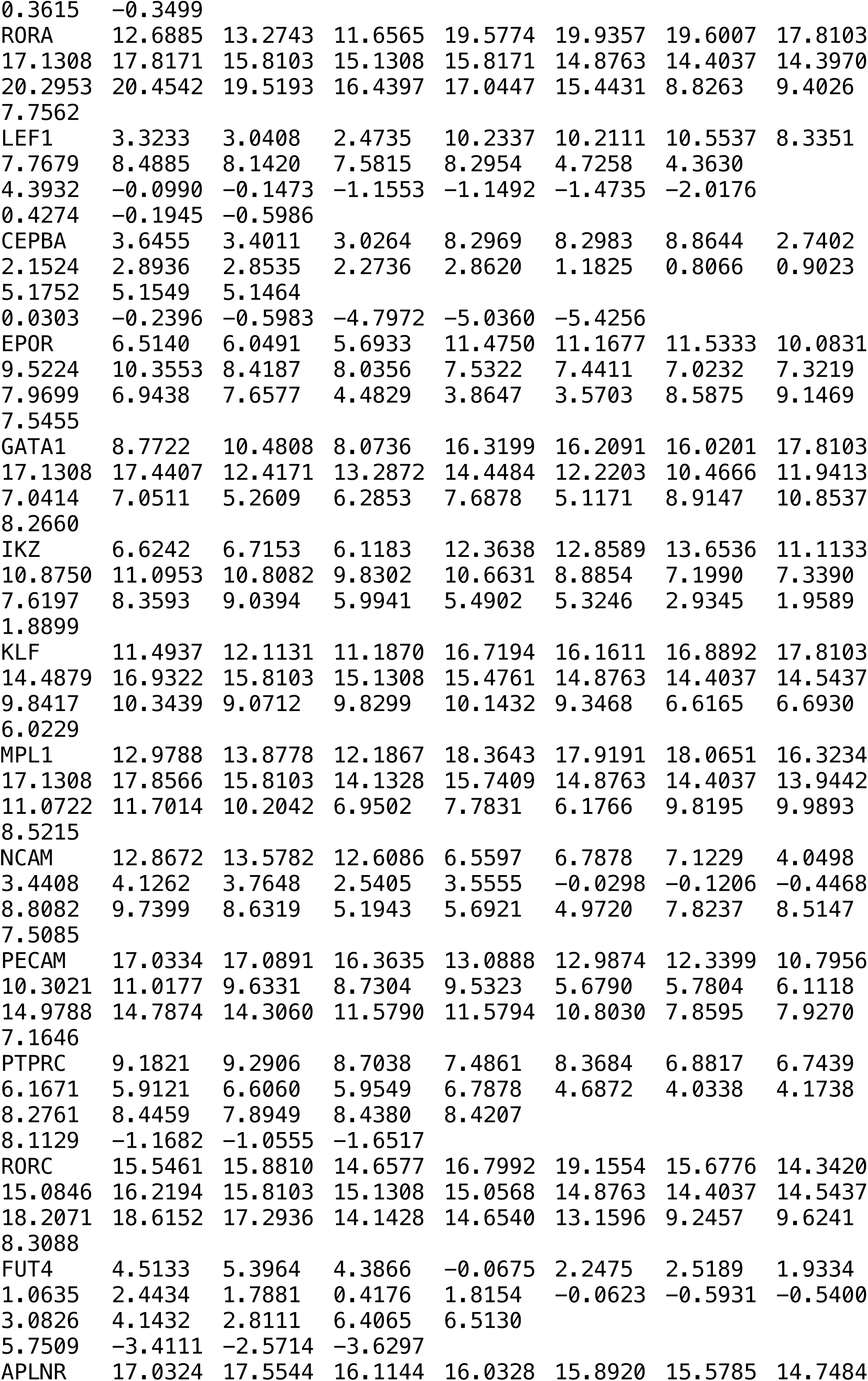
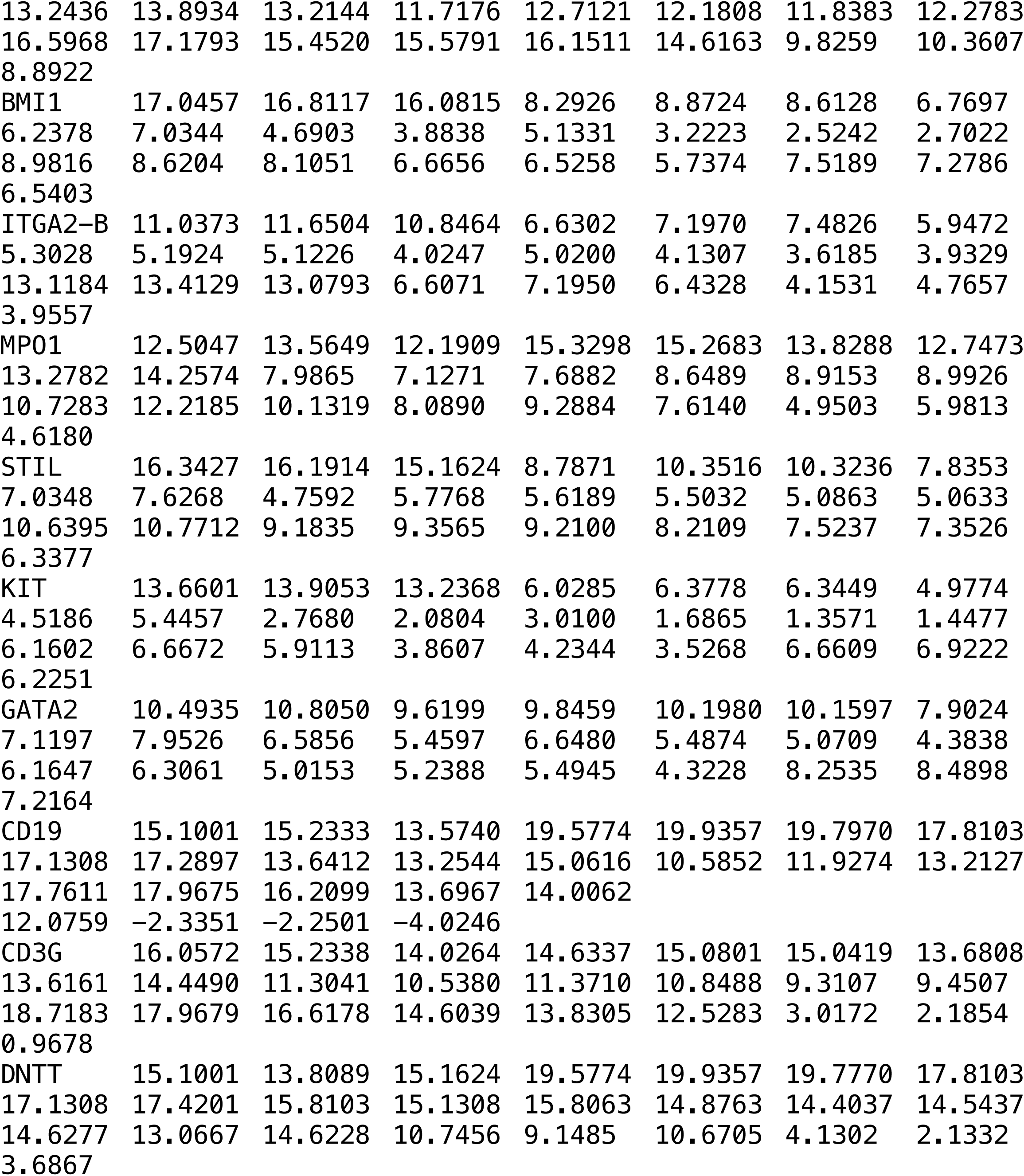
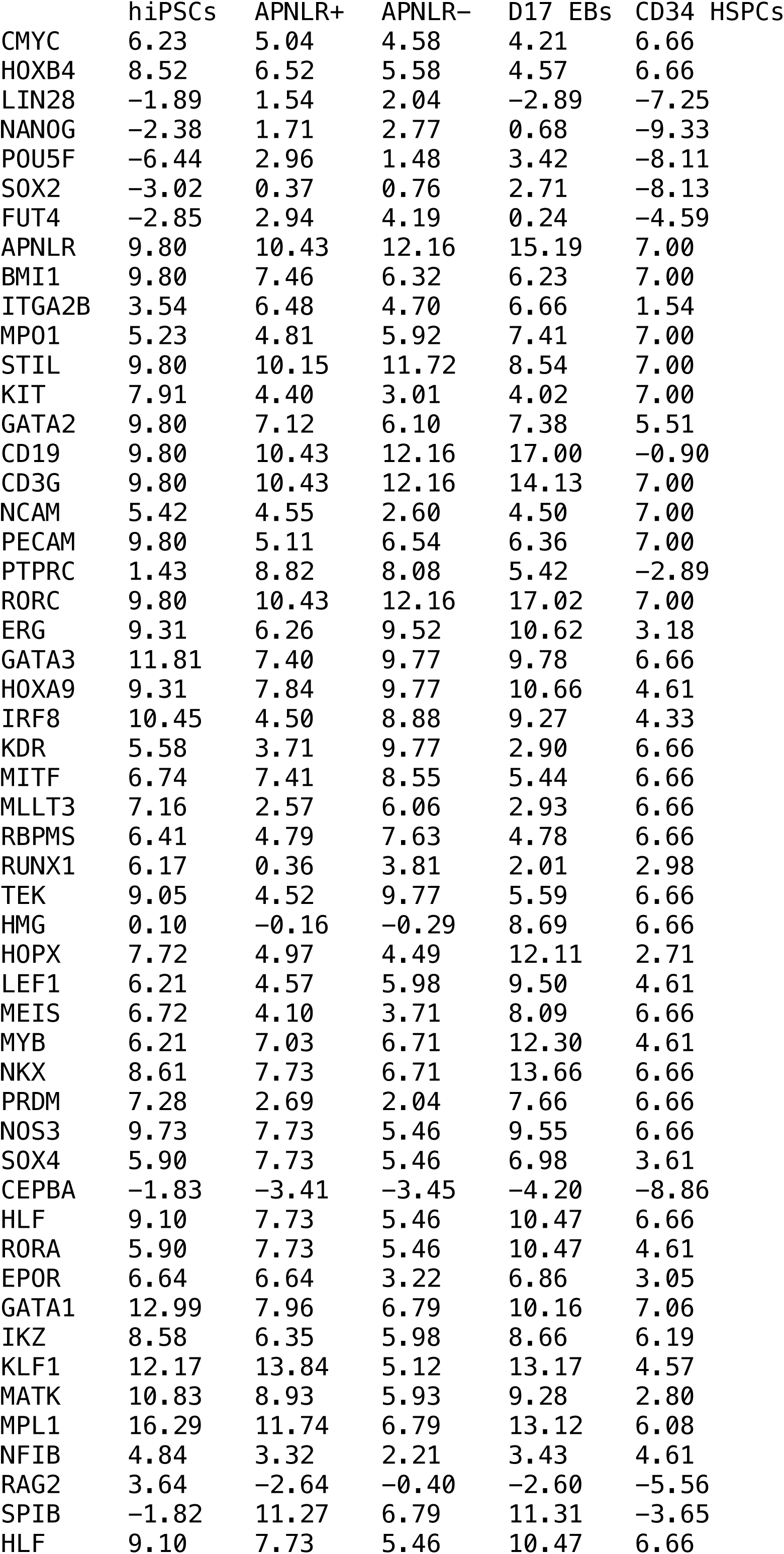
Ex *vivo* endothelial and hematopoietic potential carried by hiPSC, CD34^+^ CB HSPC and APLNR^+^, APLNR^-^ and D17 EBs. Refer to the attached supptable3.xlsx file.

